# A Cryptic Site of Vulnerability on the Receptor Binding Domain of the SARS-CoV-2 Spike Glycoprotein

**DOI:** 10.1101/2020.03.15.992883

**Authors:** M. Gordon Joyce, Rajeshwer S. Sankhala, Wei-Hung Chen, Misook Choe, Hongjun Bai, Agnes Hajduczki, Lianying Yan, Spencer L. Sterling, Caroline E. Peterson, Ethan C. Green, Clayton Smith, Natalia de Val, Mihret Amare, Paul Scott, Eric D. Laing, Christopher C. Broder, Morgane Rolland, Nelson L. Michael, Kayvon Modjarrad

## Abstract

SARS-CoV-2 is a zoonotic virus that has caused a pandemic of severe respiratory disease—COVID-19— within several months of its initial identification. Comparable to the first SARS-CoV, this novel coronavirus’s surface Spike (S) glycoprotein mediates cell entry via the human ACE-2 receptor, and, thus, is the principal target for the development of vaccines and immunotherapeutics. Molecular information on the SARS-CoV-2 S glycoprotein remains limited. Here we report the crystal structure of the SARS-CoV-2 S receptor-binding-domain (RBD) at a the highest resolution to date, of 1.95 Å. We identified a set of SARS-reactive monoclonal antibodies with cross-reactivity to SARS-CoV-2 RBD and other betacoronavirus S glycoproteins. One of these antibodies, CR3022, was previously shown to synergize with antibodies that target the ACE-2 binding site on the SARS-CoV RBD and reduce viral escape capacity. We determined the structure of CR3022, in complex with the SARS-CoV-2 RBD, and defined a broadly reactive epitope that is highly conserved across betacoronaviruses. This epitope is inaccessible in the “closed” prefusion S structure, but is accessible in “open” conformations. This first-ever resolution of a human antibody in complex with SARS-CoV-2 and the broad reactivity of this set of antibodies to a conserved betacoronavirus epitope will allow antigenic assessment of vaccine candidates, and provide a framework for accelerated vaccine, immunotherapeutic and diagnostic strategies against SARS-CoV-2 and related betacoronaviruses.

**HIGHLIGHTS:** High resolution structure of the SARS-CoV-2 Receptor-Binding-Domain (RBD).

Recognition of the SARS-CoV-2 RBD by SARS-CoV antibodies.

Structure of the SARS-COV-2 RBD in complex with antibody CR3022.

Identification of a cryptic site of vulnerability on the SARS-CoV-2 Spike.

## INTRODUCTION

The emergence of SARS-CoV-2 marks the seventh coronavirus to be isolated from humans, and the third to cause a severe disease—named COVID-19—after severe acute respiratory syndrome (SARS) and Middle East respiratory syndrome (MERS)(Munster et al., 2020). The rapid spread of SARS-CoV-2, and the grave risk it poses to global health, prompted the World Health Organization to declare, on 30 January 2020, the COVID-19 outbreak to be a public health emergency of international concern and on 11 March 2020 to be a pandemic(Wu et al., 2020; Zhou et al., 2020). As of 13 March 2020, there have been nearly 140000 SARS-CoV-2 infections and more than 5000 associated deaths reported across at least 100 countries. The rapidly evolving epidemiology of the pandemic and absence of licensed prophylactics or therapeutics for the disease have accelerated the need to elucidate the molecular biology of this novel coronavirus.

Although SARS-CoV-2 is a newly identified virus, it shares genetic and morphologic features with others in the *Coronaviridae* family, particularly those from the Betacoronavirus genus. The genome of the recently isolated SARS-CoV-2 shares 82% nucleotide identity with human SARS-CoV and 89% with bat SARS-like-CoVZXC21 (Lu et al., 2020). The spike (S) glycoprotein, in particular, bears significant structural homology with SARS-CoV compared to other coronaviruses such as MERS-CoV. Like SARS-CoV, the surface Spike (S) glycoprotein of SARS-CoV-2 binds the same host receptor, ACE-2, to mediate cell entry (Letko et al., 2020; Yan et al., 2020a). S—a class I fusion protein—is also a critical determinant of viral host range and tissue tropism and the primary target of the host immune response (Li, 2016). As such, most coronavirus vaccine candidates are based on S or one of its sub-components. Coronavirus S glycoproteins contain three segments: a large ectodomain, a single-pass transmembrane anchor and a short intracellular tail. The ectodomain consists of a receptor-binding subunit, S1, which contains two sub-domains: one at the N-terminus and the other at the C-terminus. The latter comprises the receptor-binding domain (RBD), which serves the vital function of attaching the virus to the host receptor and triggering a conformational change in the protein that results in fusion with the host cell membrane through the S2 subunit.

Recently, the molecular structure of recombinant full-length SARS-CoV-2 Spike protein was solved in a stabilized pre-fusion state, by single particle cryo-Electron Microscopy (cryo-EM), at a resolution of 3.8 Å(Wrapp et al., 2020). Despite the comprehensive structural characterization of the spike protein as a whole, movement of the RBD between “up” and “down” conformational states prevented complete modeling of the RBD domains. Subsequent cryo-EM investigations of SARS-CoV-2 provided more detail of RBD, particularly at sites that contact the human ACE-2 receptor (Yan et al., 2020a). Here, we report the first high resolution—less than 2 Å—SARS-CoV-2 RBD. Additionally, we present the antigenicity of this recombinant RBD, particularly of interest, given the equipoise in the literature regarding the binding affinities of SARS-CoV antibodies for SARS-CoV-2 RBD. Early reports, have described that the human SARS-CoV antibody, CR3022, is able to bind to the SARS-CoV-2 RBD. In the present study, we verify binding, and subsequently solved the structure of SARS-CoV-2 RBD in complex with CR3022 with a novel “cryptic” epitope.

## RESULTS

### High resolution structure of the SARS-CoV-2 RBD

The SARS-CoV-2 RBD (residues 313-532), with a C-terminal His-tag, was expressed in 293F cells, and purified by NiNTA affinity, and size-exclusion chromatography. Crystallization condition screening identified 20% Jeffamine D2000, 10% Jeffamine M2005, 0.2 M NaCl, 0.1M MES pH 5.5 for diffraction quality crystal growth. Crystals diffracted to <1.8 Å in group P 4_1_ 2_1_ 2 and to a complete dataset to 1.95 Å that could be scaled and processed (**Table 1**). The structure was refined to an R_free_ of 20% and R_work_ of 22% with no Ramachandran outliers. S residues 313-532 were clearly interpretable from the electron density map, with a dual conformation of a loop containing residues 484 to 487 clearly visible in the electron density map (**Figure 1**). Structure comparison of the unliganded RBD structure presented here, with the stabilized prefusion SARS-CoV-2 Spike (S-2P) molecule structure determined by Cryo-EM (PDB ID: 6VSB) (Wrapp et al., 2020) shows high structural similarity, with an RMSD of 0.68, 0.68, and 0.71 for each of the spike protomers. In the structure of the S-2P molecule (S-2P) (Wrapp et al., 2020) 25, 29 or 49 amino acids (aa) within each protomer RBD are not modeled, including 40% of the ACE-2 receptor binding site as measured by buried surface area (BSA)(Yan et al., 2020b). The SARS-CoV RBD-2 compared to liganded (PDB ID: 2AJF) and unliganded (PDB ID: 2GHV) SARS-CoV RBD structures shows high structural similarity, except for residues 473-488 (**Figure 1**). A chimeric SARS-CoV-2 RBD structure (PDB ID: 6VW1) with 23 aa differences compared to SARS-CoV-2, in complex with human ACE-2 was recently released in the PDB. Comparisons with previously published structures with the SARS-CoV-2 RBD highlight residues 473-488 as an area with significant structural plasticity. In the RBD structure from this study, we observe electron density for two conformations of the 482-486 loop. One of these conformations is highly similar to the ligand bound form of the RBD, while the second conformation would clash with the ACE2 receptor. Both this structural detail of the unliganded RBD and comparison to previously described RBD structures indicates that this area of the RBD is structurally malleable with implications for antibody or small molecule therapeutics design.

**Table 1.**
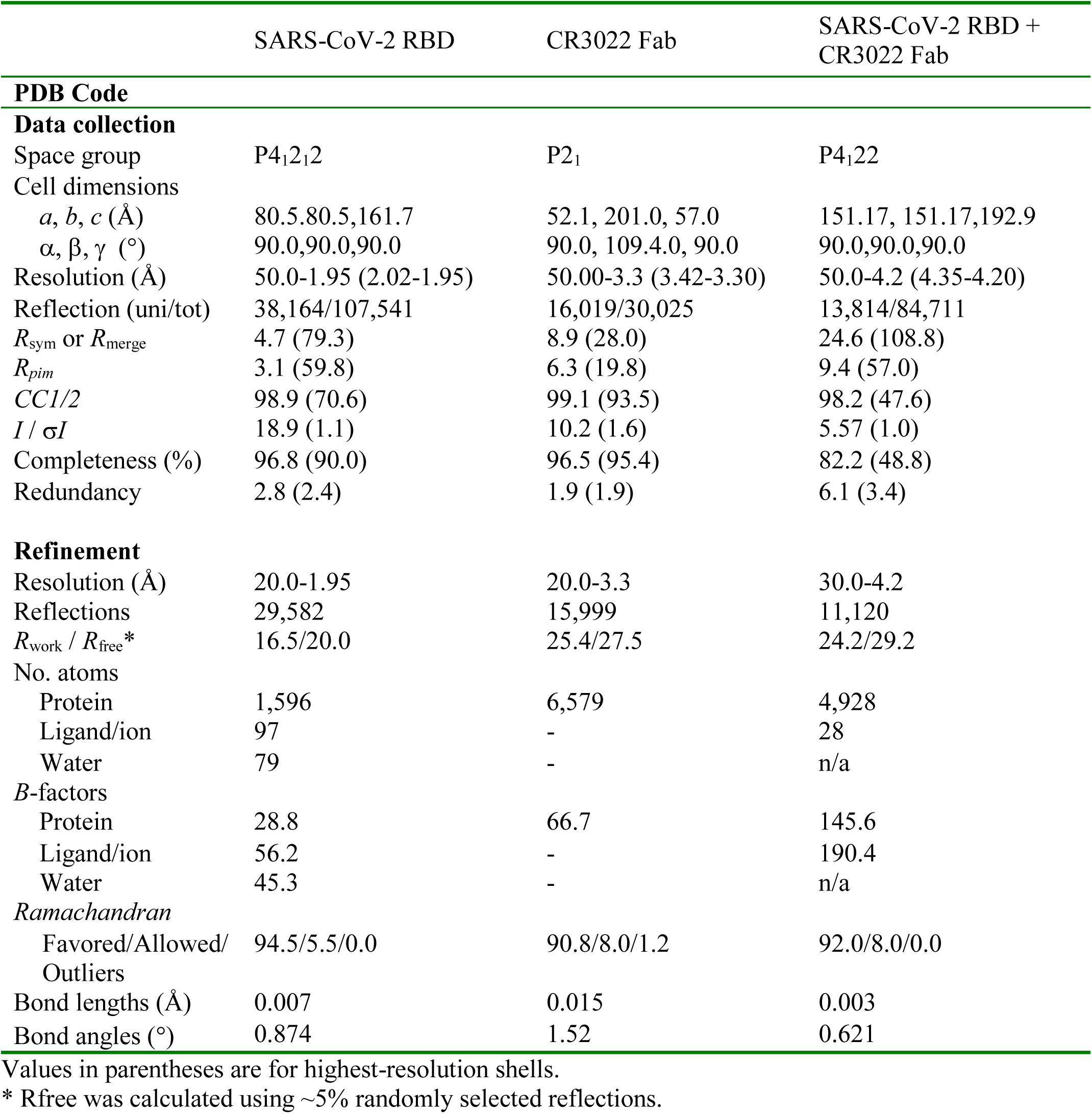
Crystallographic Data Collection and Refinement Statistics

**Figure 1.**
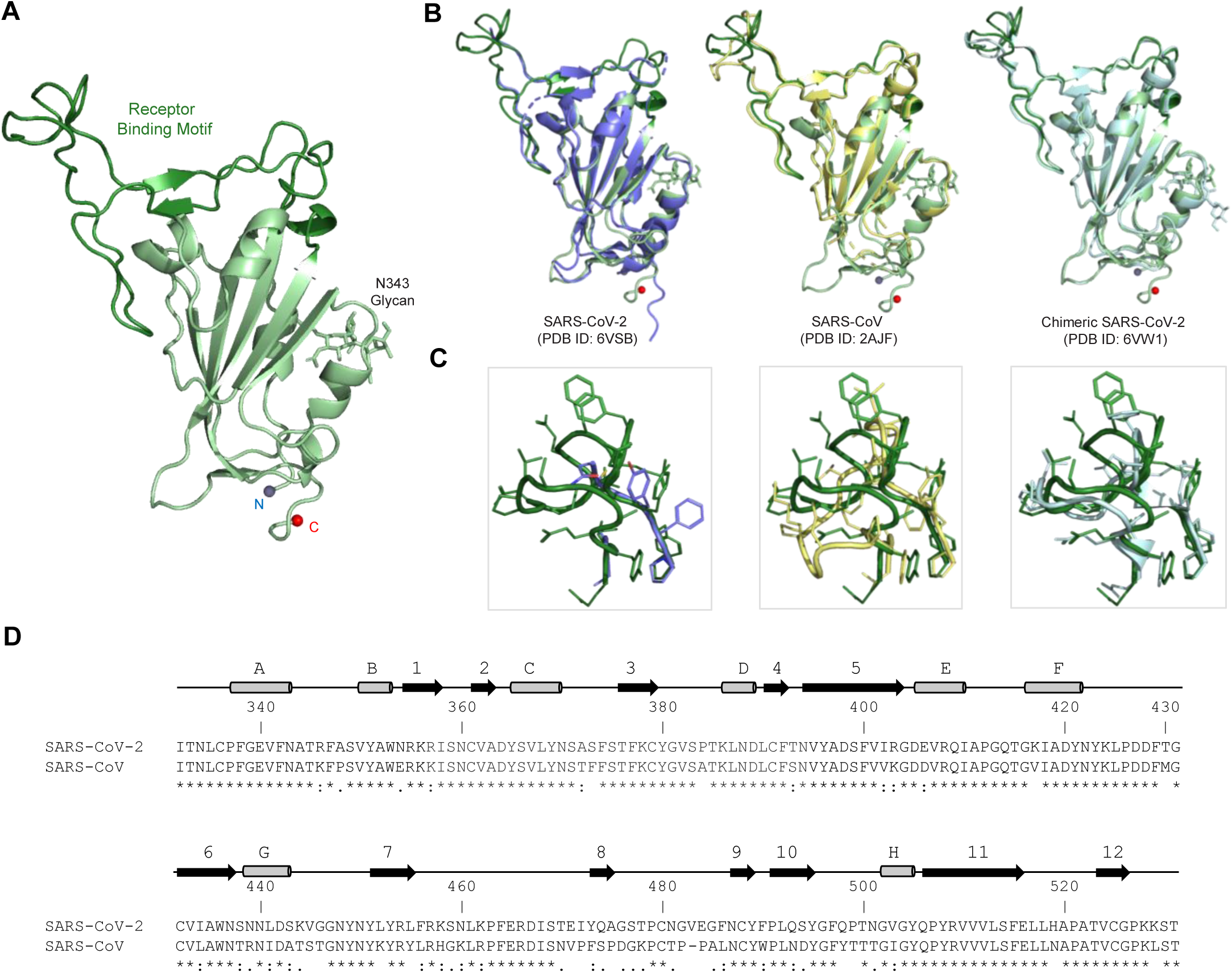
Crystal structure of the SARS-CoV-2 Receptor Binding Domain (RBD). **A** The SARS-CoV-2 RBD is shown in ribbon representation, glycan N343 is shown in stick representation, with the N- and C-termini shown as spheres. The Receptor-binding motif (residues 437-507) is colored forest green. **B** The SARS-CoV-2 RBD structure is overlaid with related structures including the incomplete RBD structure taken from the S-2P trimer structure (PDB ID: 6VSB), SARS-CoV RBD (PDB ID: 2AJF), and a chimeric SARS-CoV-2 RBD structure (PDB ID: 6VW1). **C** Close-up view of the membrane distal region of the RBD (residues 471-491). **D** Sequence alignment of SARS-CoV-2 and SARS-CoV RBDs. The RBD secondary structure is displayed above the sequence alignment Residues with significant structural difference > 2 Å are highlighted in purple.

### Identification of a set of cross-reactive SARS-CoV-2 antibodies

In an effort to identify antibodies that could bind to SARS-CoV-2, we screened a set of SARS-CoV,(Tripp et al., 2005) and MERS CoV(Wang et al., 2018; Wang et al., 2015) RBD-reactive antibodies for binding to the SARS-CoV-2 RBD. We demonstrated that the SARS-CoV mouse antibody 240CD (Tripp et al., 2005) had nanomolar (nM) affinity for the SARS-CoV-2 RBD and did not significantly block ACE-2 receptor binding (**Figure 2**). CR3022—a SARS-CoV neutralizing antibody (Tian et al., 2020) identified from a human phage-display library (ter Meulen et al., 2006)—also bound to SARS-CoV-2 RBD with nM affinity (**Figure 2B**). We assessed competition binding between 240CD and CR3022, and show that these antibodies cross-compete with each other for binding to the SARS-CoV-2 RBD (**Figure 2C, 2D**).

**Figure 2.**
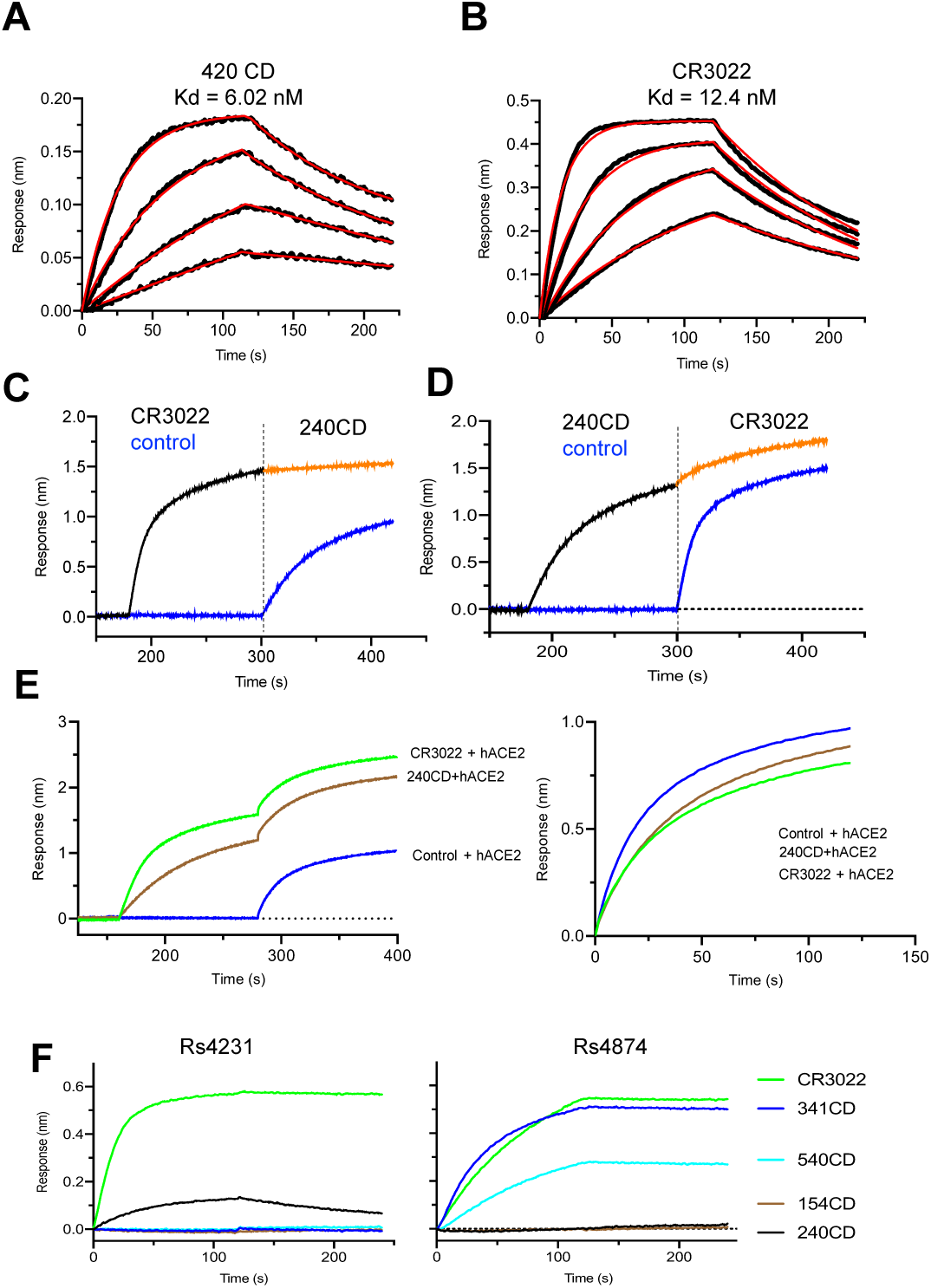
Antigenic characterization of SARS-CoV-2 RBD. **A**,**B**, Binding kinetics of 240CD and CR3022 to SARS-CoV-2 RBD measured by biolayer interferometry. Kinetic constants were determined were calculated using a minimum of four dilutions of the RBD and fitted using a 1:1 binding model. **C**,**D**, Competition binding of antibodies CR3022 and 240CD to SARS-CoV-2 RBD. CR3022 or control antibody was allowed to bind to SARS-COV-2 prior to binding to 240CD or vice-versa. **E** SARS-CoV-2 RBD was sequentially bound by antibodies CR3022 or 240CD followed by soluble human ACE2 receptor. **F** SARS-CoV reactive antibodies were assessed for binding to bat SARS-related CoV Rs4784 and Rs4231 S glycoproteins.

SARS-CoV-2 has a likely zoonotic origin and horseshoe bats have been implicated as natural reservoirs of both SARS-CoV and SARS-CoV-2 (Menachery et al., 2015; Zhou et al., 2020). As such, we next explored antibody cross-reactivity with the S glycoproteins of two bat SARS-related CoVs: SARSr-CoV Rs4874 (Ge et al., 2013; Yang et al., 2015) and Rs4231 (Hu et al., 2017), which are closely related to the progenitor of SARS-CoV and retain the ability to utilize human ACE-2. CR3022 was able to recognize a recombinant Spike glycoproteins generated from bat SARSr-CoV Rs4874, while 240CD, and other mouse generated monoclonal antibodies have a mixed recognition phenotype (**Figure 2F**).

### Crystal structure of antibody CR3022 in complex with SARS-CoV-2 RBD

The antigenic cross-reactivity of this set of antibodies precipitated an investigation into their molecular recognition determinants. The potential relevance of a human antibody motivated the investigation to prioritize studies of CR3022, for which a sequence was available (ter Meulen et al., 2006). The CR3022 heavy chain is encoded by IGHV5-51*03, contains a 12-aa CDR H3 with 8 V gene-encoded residues altered by somatic hypermutation. CR3022 light chain is encoded by IGKV4-1*01 with 1 V gene-encoded residue, altered by somatic hypermutation, and a 9-aa CDR L3 (**Extended Data Figure 2A**). To provide an atomic-level understanding of the structure of the CR3022 antibody, we crystallized the antigen-binding fragment (Fab) of CR3022. Crystals diffracted to 3.2 Å resolution in space group P 2_1_ (**Table 1**). Overall the structure of the CR3022 Fab revealed a relatively flat antigen-combining site, with the exception of an extended protruding 12-aa CDR L1 loop (**Figure S2A**).

To determine the structure of CR3022 in complex with the SARS-CoV-2 RBD, we carried out crystallization conditions screening, with crystals of the CR3022-RBD complex forming in 1M Succinic acid, 0.1M Hepes pH 7, 2% PEG MME2000 and determined the crystal structure by X-ray diffraction to 4.25 Å (**Table 1**). The complex structure was solved by molecular replacement using the refined CR3022 and SARS-CoV-2 RBD structures as search models and was refined to an R_work_/R_free_ of 0.242/0.292 (**Table 1**). CR3022 bound to the RBD at an epitope centered on S glycoprotein residues 377-386 with a total buried surface area of 871 Å (**Figure 3, Figure S2**, and **Table S1**). This region is highly conserved between SARS-CoV and SARS-CoV-2 (**Figure S3**). Comparison of the CR3022 epitope site with previously described antibody-complex structures for SARS-CoV, and MERS-CoV indicates that CR3022 describes a novel recognition site (**Figure 3** and **Figs. S3, S4**). Further sequence analysis of the epitope indicates that this epitope is conserved in betacoronavirus clade 2b, with also some similarity in clade 2d (**Fig. S3**). To confirm that this site was also shared with 240CD, we produced an RBD knockout mutant by introducing a glycan sequon at position 384, and by biolayer interferometry show that both CR3022 and 240CD binding to the RBD can be eliminated by the introduction of a glycan at this site (**Fig. S2D**).

**Figure 3.**
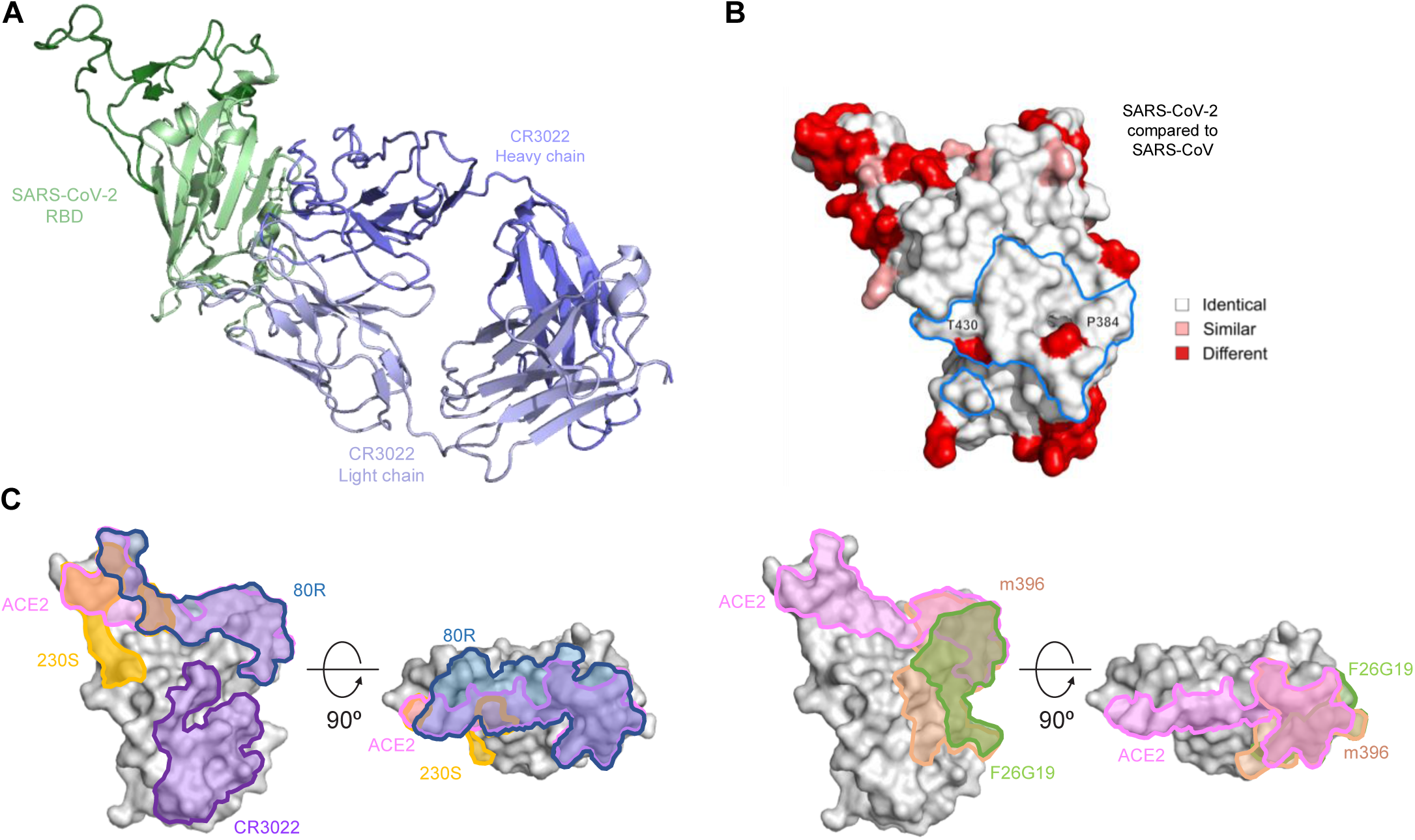
Crystal structure of CR3022 in complex with SARS-CoV-2 RBD. **A**, RBD and CR3022 are shown in cartoon representation. **B** Structure of the SARS-CoV-2 RBD shown in surface representation. Residues which differ between SARS-CoV-2 and SARS-CoV are colored red. The CR3022 epitope is outlined in blue, and Thr-430 and Phe-384 located within the epitope are labeled. **C** The location of antibody CR3022, 230S, 80R, m396 and F26G19 epitopes and ACE-2 binding site on the RBD are outlined on the surface of the SARS-CoV-2 RBD.

### Identification of a cryptic site of vulnerability recognized by CR3022

The epitope conservation within the clade explains the antigenic cross-reactivity with both human SARS-CoV and bat SARS related CoV. To date, there has been extensive structural characterization of the SARS-CoV, and MERS-CoV spike molecule and domains, which provides a framework for understanding the novel SARS-CoV-2 spike molecule (**Table S2**). In the context of the coronavirus trimeric S glycoproteins, the RBD displays two prototypical conformations either in an “up” or “down” position, with implications for receptor binding and cell entry (**Table S1**). To further analyze these conformations, we modeled the CR3022 binding to the trimeric structures of SARS-CoV-2, SARS-CoV and MERS-CoV. The CR3022 epitope is occluded by adjacent spike protomers when the RBD is in the “down” conformation, but becomes more accessible when the spike is in a more open conformation here multiple RBD molecules are in the “up” conformation (**Figure 4**). There is still a clash of the antibody Fc1 region with the NTD from the same protomer, or an RBD from an adjacent protomer when modeled using the static structure.

**Figure 4.**
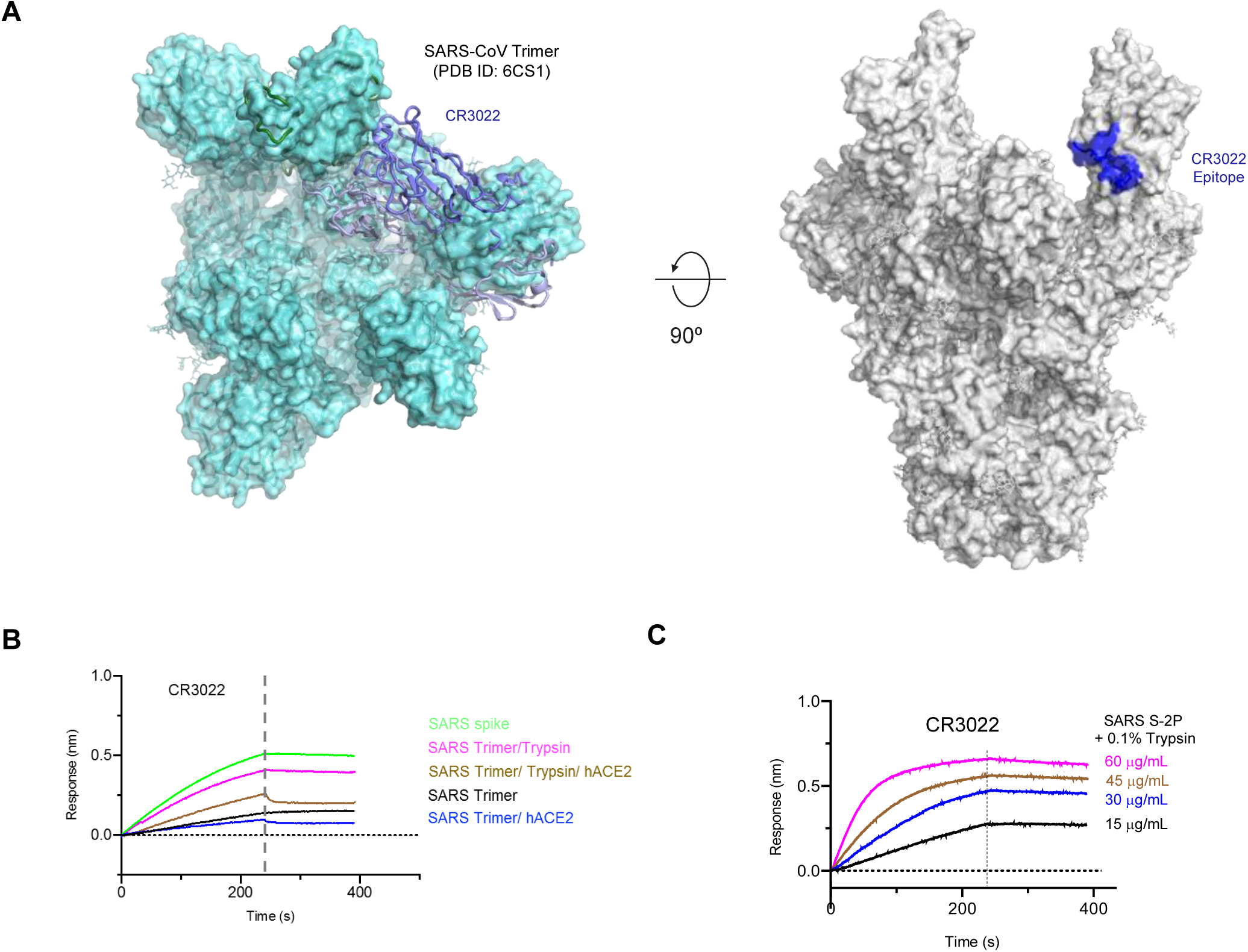
Identification of CR3022 epitope as a “cryptic” epitope. Structural alignment of the SARS-CoV-2 RBD-CR3022 complex with the SARS-CoV S-2P structure. **A** The RBD-CR3022 structure is aligned to the SARS-CoV trimer structure (surface representation; PDB ID: 6CS1), where two RBD molecules are located in the “up” conformation. In this static structure, the Fc1 region of CR3022 (ribbon representation) clashes with the NTD of the same protomer. However, the epitope is fully accessible when more than one RBD is in the “up” representation. **B** Biolayer interferometry measurement of CR3022 binding to SARS S proteins with trypsin treatment or ACE2 receptor binding. **C** CR3022 binding to a serial dilution of SARS S-2P protein following trypsin treatment.

To understand whether CR3022 could bind to SARS-CoV S glycoproteins, we measured binding to stabilized S-2P or non-stabilized versions of S (**Figure 4**). We observed robust binding to the non-stabilized S glycoprotein, while binding to SARS S-2P Trimer was low. We then treated the SARS S-2P trimer with trypsin and/or incubation with the ACE2 receptor to assess whether minimal proteolytic action or receptor binding could increase the availability of the “cryptic” CR3022 epitope. Incubation of the stabilized S-2P trimer with human ACE2 did not dramatically affect CR3022 binding, while in contrast, the trypsin treatment of the S-2P protein resulted in increased binding akin to the unstabilized S glycoprotein binding, and the level of binding was titratable, with increasing amounts of S-2P resulting in higher CR3022 binding. Given the prior neutralization and protection studies utilizing CR3022, and its ability to complement potent neutralizing antibodies, it is likely that the CR3022 epitope represents a “cryptic” epitope that becomes exposed during the processes of viral cell entry.

In summary, our data represents the most detailed structural information for the SARS-CoV-2 RBD to date and the first structure of the SARS-CoV-2 in complex with a human antibody. The presence of “cryptic” but protective epitopes for influenza (Bangaru et al., 2019), and Ebola viruses (West et al., 2018), have been previously described. The identification of a novel “cryptic” epitope for betacoronaviruses including SARS-CoV, and SARS-CoV-2 highlight a novel viral vulnerability that can be harnessed in combination with ACE2 receptor site targeting monoclonal antibodies for vaccine and therapeutic countermeasure development.

## ACKNOWLEDGEMENTS

This work was supported by funding from the Defense Health Agency, as well as a cooperative agreement (W81XWH-07-2-0067) between the Henry M. Jackson Foundation for the Advancement of Military Medicine, Inc., and the U.S. Department of Defense (DoD). Funding from Biological Defense Research Directorate of the Naval Medical Research Center (HT9404-13-1-0021; Component Project: Soluble Trimeric Filovirus Envelope Glycoproteins) and Defense Threat Reduction Agency (HDTRA1-17-C-0019; Chulalongkorn Luminex Training and Research Preparedness) to CCB. X-ray diffraction data were collected at beamlines ID-24-C and ID-24-E at the Advanced Photon Source, Argonne National Laboratory. The Northeastern Collaborative Access Team (NE-CAT) beamlines are funded by the National Institute of General Medical Sciences from the National Institutes of Health (P41 GM103403) at the Advanced Photon Source, Argonne National Laboratory. Reagents were obtained through BEI Resources, NIAID, NIH: Monoclonal Anti-SARS-CoV S glycoprotein (Similar to 240C), NR-616; (Similar to 341C), NR-617; (Similar to 540C), NR-618. We thank the authors who made their SARS-CoV-2 genome sequences available through GISAID or GenBank. The opinions or assertions contained herein are the private views of the authors, and are not to be construed as official, or as reflecting true views of the Department of the Army or the Department of Defense.

## METHODS AND MATERIALS

### Production of recombinant proteins

The Shanghai Public Health Clinical Center & School of Public Health, in collaboration with the Central Hospital of Wuhan, Huazhong University of Science and Technology, the Wuhan Center for Disease Control and Prevention, the National Institute for Communicable Disease Control and Prevention, Chinese Center for Disease Control, and the University of Sydney, Sydney, Australia released the sequence of a coronavirus genome from a case of a respiratory disease from Wuhan on January 10^th^ available at recombinomics.co/topic/4351-wuhan-coronavirus-2019-ncov-sequences/. The sequence was also deposited in GenBank (accession MN908947) and GISAID (>EPI_ISL_402125). DNA encoding the SARS-Cov-2 RBD (residues 331-527) was synthesized (Genscript) with a C-terminal His6 purification tag and cloned into a CMVR plasmid, and protein was expressed by transient transfection in 293F cells for six days. The SARS-CoV-2 RBD-His protein was purified from cell culture supernatant using a Ni-NTA (Qiagen) affinity column. DNA encoding the S protein ectodomains (residues 1-1194) from bat SARS-related CoV isolates Rs4231 and Rs4874 (ref.(Hu et al., 2017)) were synthesized (Genscript) with a C-terminal T4-Foldon domain or C-terminal GCN domain, respectively, followed by factor xA cleavage sites and Strep-Tactin purification tags. Bat SARSr-CoV S genes were cloned into a modified pcDNA3.1 expression plasmid (Chan et al., 2009). Protein was initially expressed by transient transfection in 293F cells for six days, then serial cloned to select stably expressing cell lines (Yan L., in submission). The Rs4231-T4 and Rs4874-GCN S proteins were purified from cell culture supernatant using a Strep-Tactin affinity column. The oligomeric structure of these S proteins was selected by size exclusion chromatography (GE/AKTA) and trimeric S proteins were confirmed by Native-PAGE. SARS S-2P was produced as previously described, with Strep-Tactin affinity chromatography followed by gel filtration using a 16/60 Superdex-200 purification column. Purification purity for all S glycoproteins was assessed by SDS-PAGE.

The sequences of the CR3022 variable regions of the heavy and light chains are available in GenBank under accession numbers DQ168569 and DQ168570, respectively (ter Meulen et al., 2006). These sequences were synthesized (Genscript) and cloned into CMVR expression vectors (NIH AIDS reagent program) between a murine Ig leader (GenBank DQ407610) and the constant regions of human IgG1 (GenBank AAA02914), Igκ (GenBank AKL91145). Plasmids encoding heavy and light chains were co-transfected into Expi293F cells (ThermoFisher) according to the manufacturer’s instructions. After 5 days, antibodies were purified from cleared culture supernatants with Protein A agarose (ThermoFisher) using standard procedures, buffer exchanged into Phosphate-Buffered Saline (PBS), and quantified using calculated E and A280 measurements.

The Fab fragment of antibody CR3022 was prepared by digestion of the full-length IgG using enzyme Lys-C (Roche). The digestion reaction was allowed to proceed for 2.5 hours at 37°C. Digestion was assessed by SDS-PAGE and upon completion, the reaction mixture was passed through protein-G beads (0.5-1 ml beads), 3 times and the final flow through was assessed by SDS-PAGE for purity. The Fab fragment was mixed with purified SARS-CoV-2 RBD, and the complex was allowed to form for 1 hour at room temperature.

### Sequence information

SARS-CoV-2 RBD (signal peptide is underlined, purification tag in italics)

<u>MDSKGSSQKGSRLLLLLVVSNLLLPQGVVG</u>NITNLCPFGEVFNATRFASVYAWNRKRISNCVAD YSVLYNSASFSTFKCYGVSPTKLNDLCFTNVYADSFVIRGDEVRQIAPGQTGKIADYNYKLPDDF TGCVIAWNSNNLDSKVGGNYNYLYRLFRKSNLKPFERDISTEIYQAGSTPCNGVEGFNCYFPLQS YGFQPTNGVGYQPYRVVVLSFELLHAPATVCG*PGSHHHHHH*

CR3022 Heavy chain Fv

EVQLVQSGTEVKKPGESLKISCKGSGYGFITYWIGWVRQMPGKGLEWMGIIYPGDSETRYSPSFQGQVTISADKSINTAYLQWSSLKASDTAIYYCAGGSGISTPMDVWGQGTTVTVSS

CR3022 Light chain Fv

DIVMTQSPDSLAVSLGERATINCKSSQSVLYSSINKNYLAWYQQKPGQPPKLLIYWASTRESGVPDRFSGSGSGTDFTLTISSLQAEDVAVYYCQQYYSTPYTFGQGTKVEIK

### Cell lines

Expi293F (ThermoFisher Scientific #A14527), and 293F cell lines were utilized in this study.

### X-ray Crystallography

#### Crystallization

SARS-CoV-2 RBD at 10 mg/ml and 5 mg/ml in PBS buffer was screened for crystallization conditions using an Art Robbins Gryphon crystallization robot, 0.2 ul drops, and a set of 1200 crystallization conditions. Crystal drops were observed using a Jan Scientific UVEX-PS with automated UV and brightfield drop imaging robot. Crystals of the SARS-CoV-2 RBD grew after 24 hours in multiple conditions from the Molecular Dimensions MIDAS crystal screen, with diffraction-quality crystals seen in conditions B1, G1, F6, and H10. CR3022 Fab was screened for crystallization at 10.0 mg/ml and 5.0 mg/ml concentrations in PBS. Diffraction quality crystals grew after 48 hours in 0.1M Imidazole pH 6.5, 40% 2-propanol and 15% PEG 8,000. For the complex, CR3022 Fab and SARS-CoV-2 RBD were mixed in 1:1 molar ratio and crystallization drops were set-up at 8.0 and 4.0 mg/ml concentrations in PBS buffer as described above. Crystals grew in a crystallization condition containing 1M Succinic acid, 0.1M HEPES pH 7.0 and 2% PEG MME2000. Both, RBD alone and CR3022 Fab-RBD complex, crystals were harvested and cryo-cooled in their respective crystallization conditions plus 25% glycerol.

#### Diffraction data collection and processing

Single crystals were transferred to mother liquor containing 22% glycerol, and cryo-cooled in liquid nitrogen prior to data collection. Diffraction data for SARS-CoV-2 RBD were collected at Advanced Photon Source (APS), Argonne National Laboratory, NE-CAT ID24-C beamline, and measured using a Dectris Eiger 16M PIXEL detector. Crystals grown in MIDAS condition B1 (20% Jeffamine D2000, 10% Jeffamine M2005, 0.2 M NaCl, 0.1M MES pH 5.5) provided the highest resolution diffraction with spots visible to 1.8 Å. A complete dataset could be processed to 1.95 Å in space group P41212. CR3022 Fab crystals diffracted to 3.3 Å on NE-CAT ID24-C beamline. Diffraction data could be scaled in P21 space group with 99.9% completeness. Diffraction data for CR3022 and SARS-CoV-2 RBD complex were collected on NE-CAT ID24-C beamline at Advanced Photon Source (APS), and measured using a Dectris Eiger 16M PIXEL detector. Diffraction data from multiple crystals were merged and scaled together to achieve a final resolution of 4.2 Å with overall completeness of 82.2%. Data collection statistics are reported in Table 1.

#### Structure solution and refinement

Phenix xtriage was used to analyze the scaled diffraction data produced from HKL2000 and XDS. Data was analyzed for completeness, Matthew’s coefficient, twinning or pseudo-translational pathology. The structure of the SARS-CoV-2 RBD was determined by molecular replacement using Phaser and a search model of the SARS RBD (PDB ID: 2AJF, molecule C). CR3022 Fab crystal structure was determined by molecular replacement using Coxsackievirus A6 neutralizing antibody 1D5 (PDB ID: 5XS7) as a search model. The CR3022-RBD complex structure was determined by molecular replacement using the refined CR3022 and SARS-CoV-2 RBD structures as search models. Refinement was carried out using Phenix refine with positional, global isotropic B factor refinement, and defined TLS groups, with iterative cycles of manual model building using COOT. Structure quality was assessed with MolProbity. The final refinement statistics for all the structures are reported in Table 1. All structure figures were generated using PyMOL (The PyMOL Molecular Graphics System [DeLano Scientific]).

### Structure comparisons

#### Weighing epitope sites based on antigen-antibody interactions

Epitope sites correspond to antigen sites that are in contact with the antibody in the antigen-antibody complex (i.e. all sites that have non-hydrogen atoms within 4 Å of the antibody). For a given epitope site, the weight, which characterizes the interaction between the epitope site and the antibody (improved based on (Bai et al., 2019)), was defined as: 

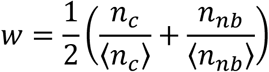

in which, *n*_*c*_ is the number of contacts with the antibody (i.e. the number of non-hydrogen antibody atoms within 4 Å of the site); *n*_*nb*_ is the number of neighboring antibody residues; ⟨*n*_*c*_⟩ is the mean number of contacts *n*_*c*_ and ⟨*n*_*nb*_⟩ is the mean number of neighboring antibody residues *n*_*nb*_ across all epitope sites. A weight of 1.0 is attributed to the average interaction across all epitope sites. Neighboring residue pairs were identified by Delaunay tetrahedralization of side-chain centers of residues (C_α_ is counted as a side chain atom, pairs further than 8.5 Å were excluded). Quickhull(Barber, 1996) was used for the tetrahedralization and Biopython PDB (Hamelryck and Manderick, 2003) to handle the protein structure. In the SARS-CoV-2 and SARS-CoV-1 RBD comparison, residues were considered similar for the following residues pairs: RK, RQ, KQ, QE, QN, ED, DN, TS, SA, VI, IL, LM and FY.

### Biolayer interferometry

Affinity kinetic interactions between SARS-CoV-2 RBD proteins and antibodies were monitored on an Octet RED96 instrument (FortéBio). After reference subtraction, binding kinetic constants were determined, from at least 4 concentrations of Fab, by fitting the curves to a 1:1 Langmuir binding model using the Data analysis software 9.0 (FortéBio). Antibodies were loaded at 30 ug/ml onto a AHC probe for 120 s followed by baseline incubation for 30-60 s.

To assess antibody competition, either 240CD or CR3022 or a non-specific control antibody CR1-07 was incubated with the SARS-CoV-2 RBD prior to assessment of binding to CR3022 or 240CD. Antibody concentration was 30 ug/ml. To assess binding of human ACE-2 receptor in the presence or absence of antibodies CR3022, or 240CD, RBD was loaded onto a HIS probe. The RBD was then sequentially incubated with either CR3022, 240CD or control antibody CR1-07 prior to incubation with human ACE-2 receptor.

CR3022 was loaded onto an AHC probe for 120s prior to incubation with SARS-CoV S glycoproteins (15 ug/ml) alone or pre-incubated with ACE2 protein. SARS S-2P protein was treated with 0.1% bovine pancreas trypsin for 10 minutes prior to binding to binding measurements. SARS Spike protein was provided by BEI resources, Lot 768P152. Binding of CR3022 was also carried out against a series of concentrations of SARS S-2P which had been treated with 0.1% w/w bovine pancreatic trypsin.

## DATA AVAILABILITY

The associated accession numbers for the coordinates and structure factors reported in this paper are being deposited to the PDB.

## ADDITIONAL RESOURCES

None

---

**Table S1.**
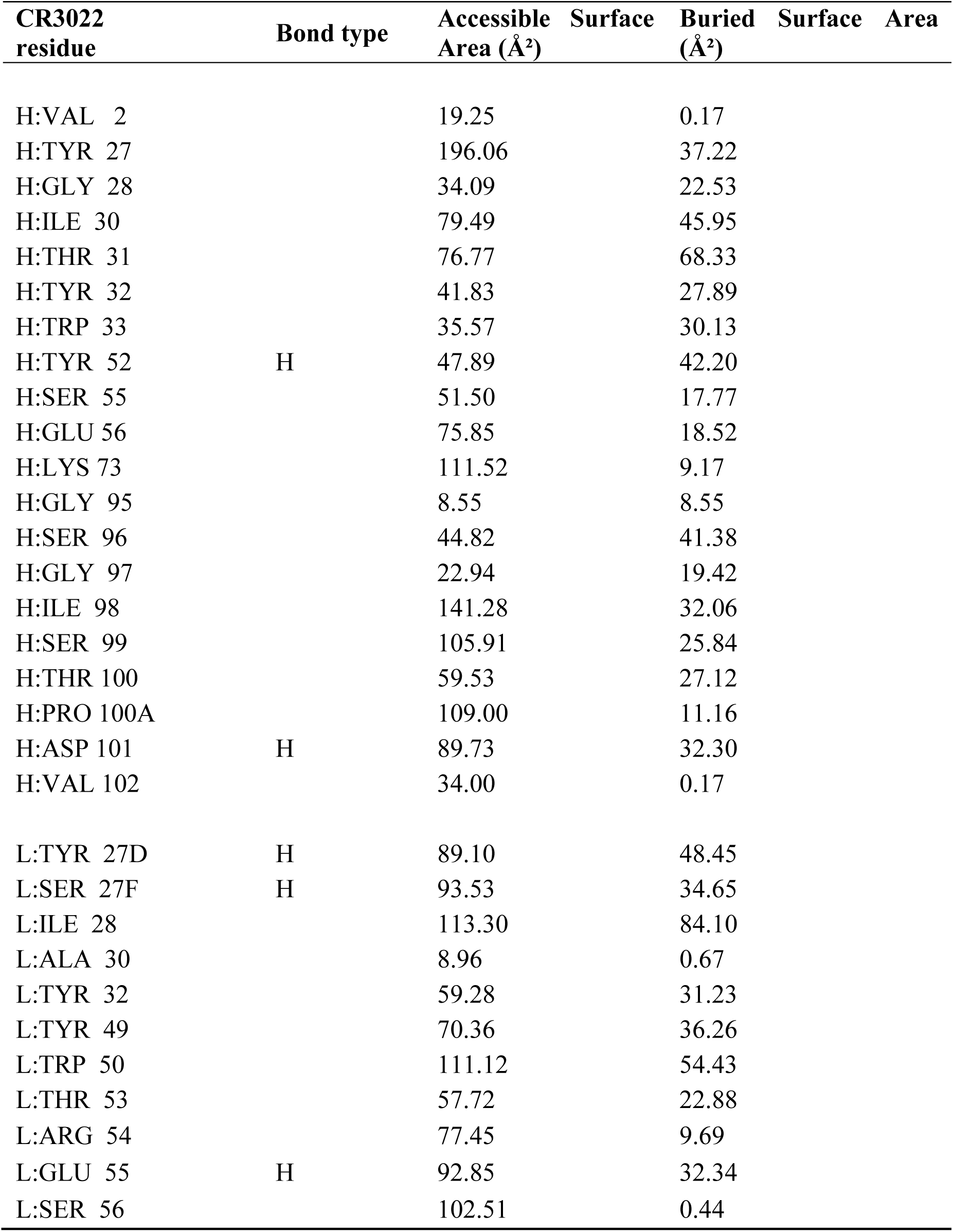
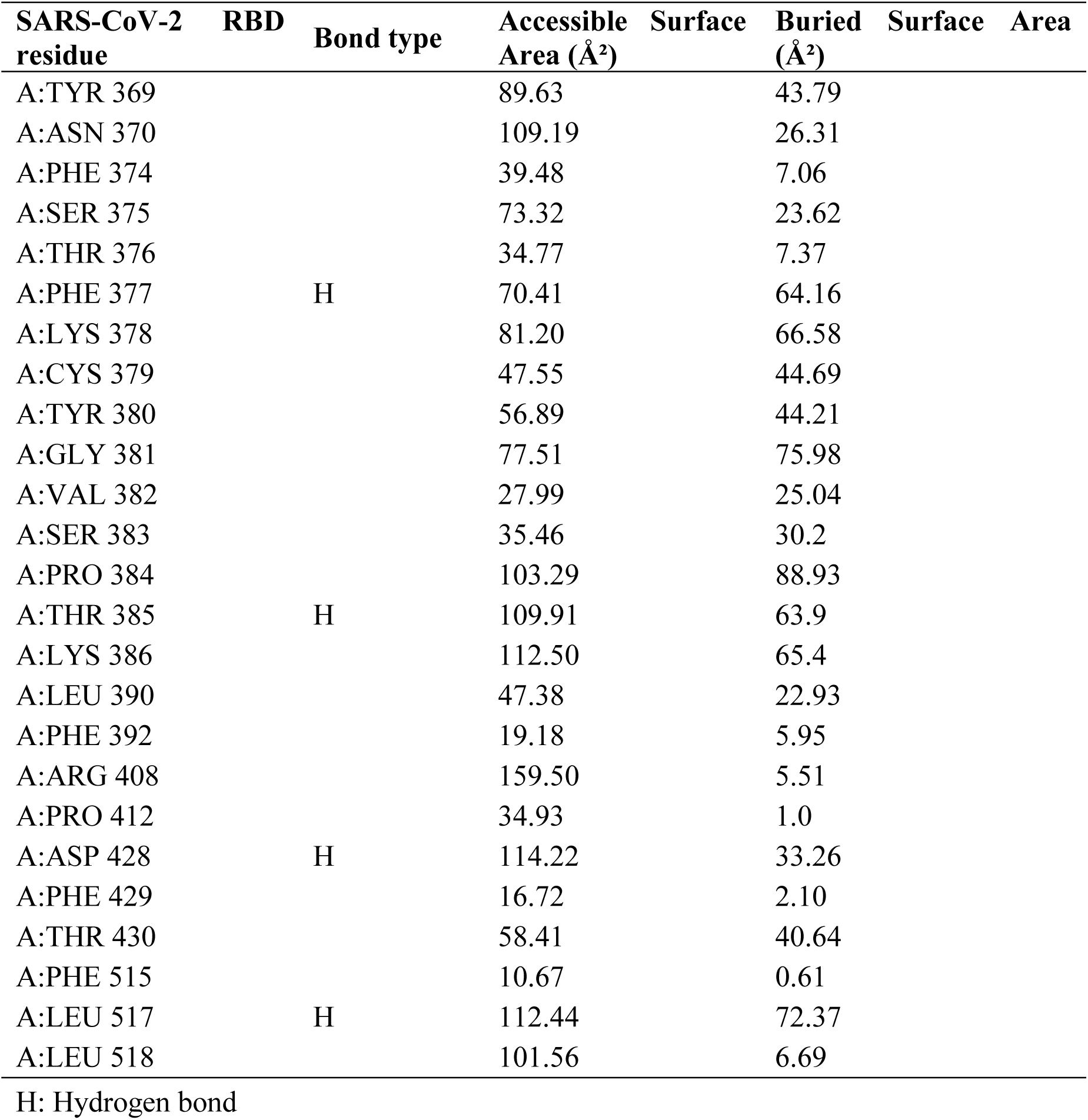
Buried surface area of CR3022 in complex with SARS-CoV-2 RBD

**Table S2.**
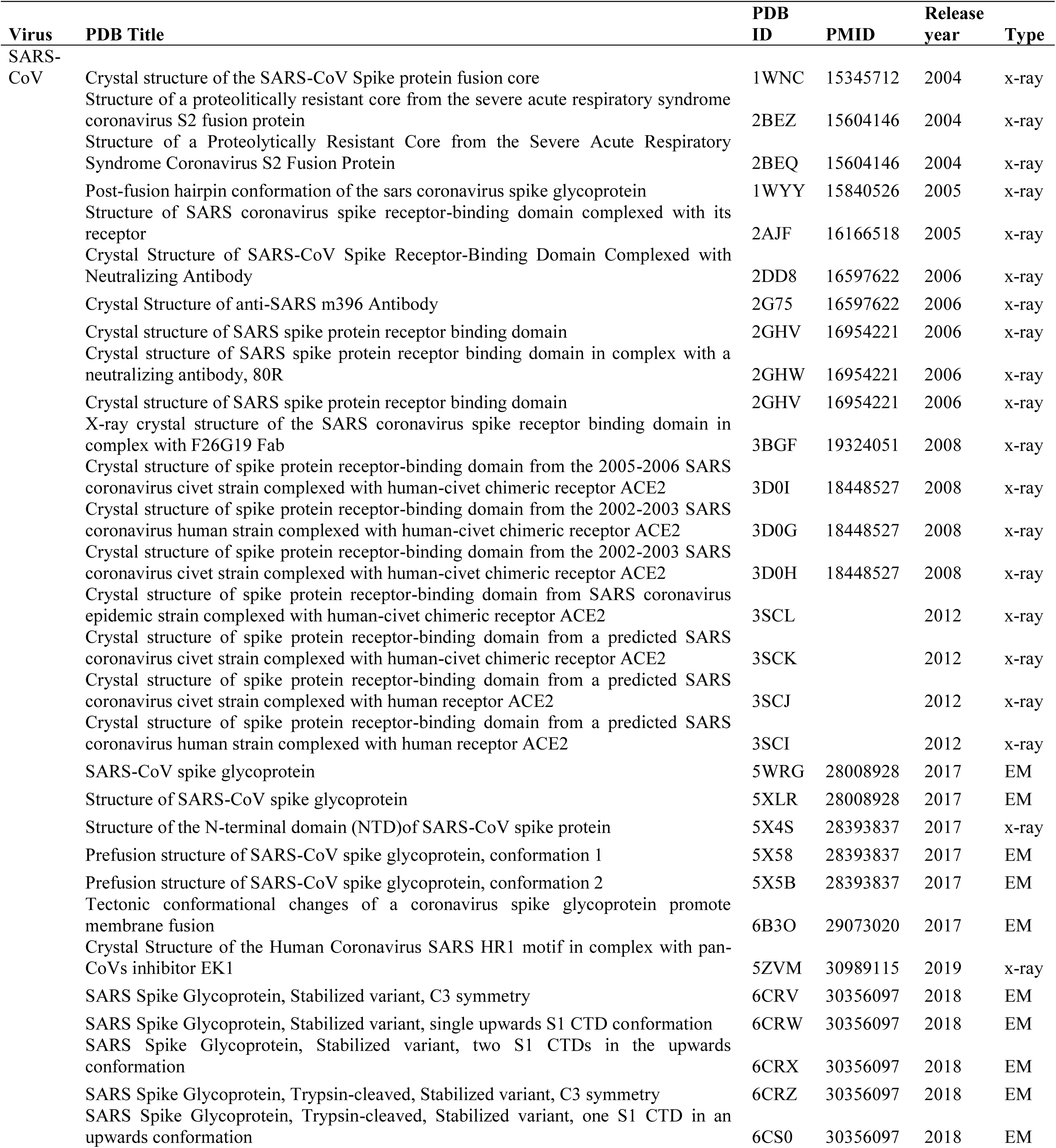
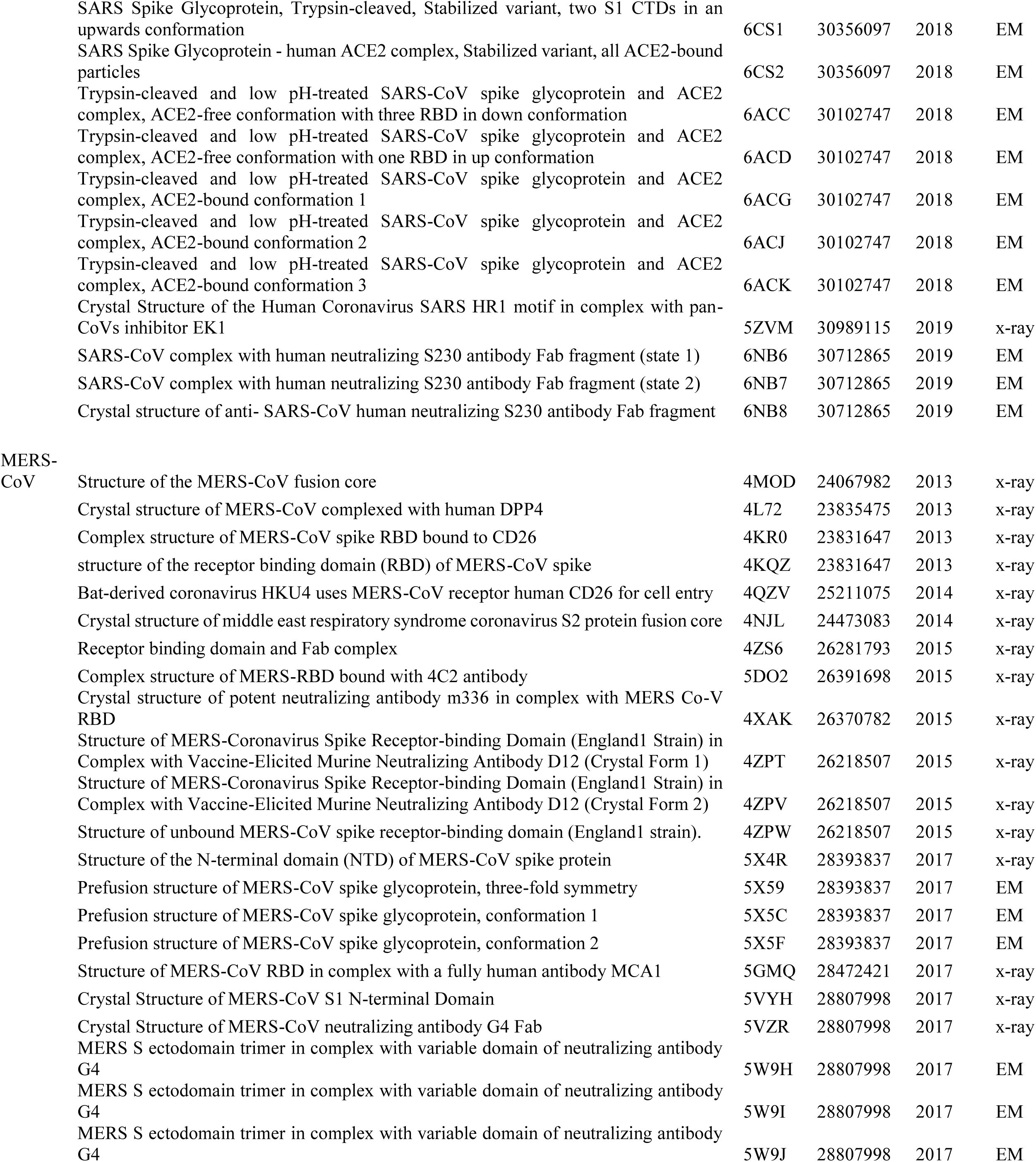
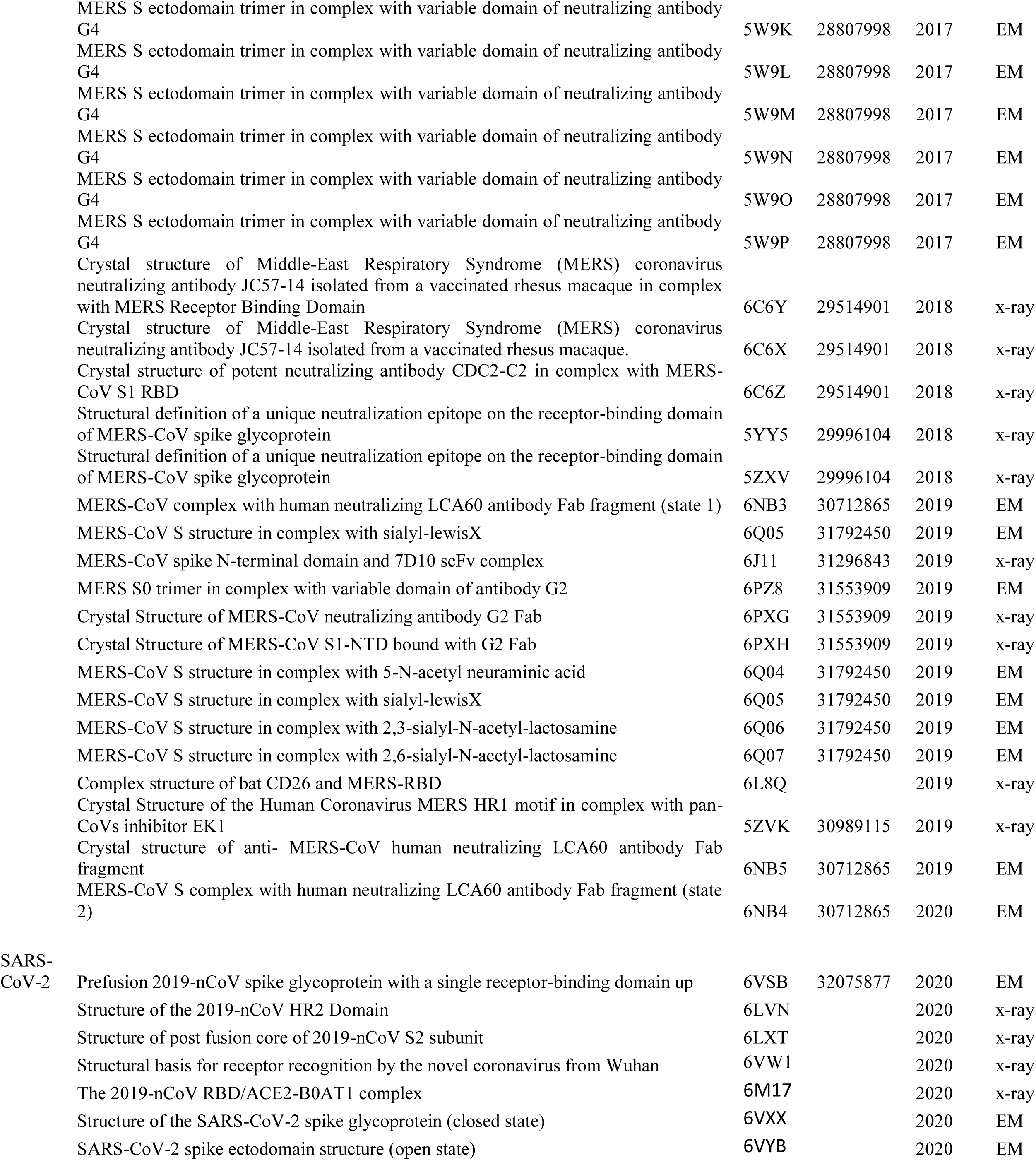
Protein DataBank files of SARS-CoV, MERS-CoV, and SARS-CoV-2 S structures

**Fig. S1.**
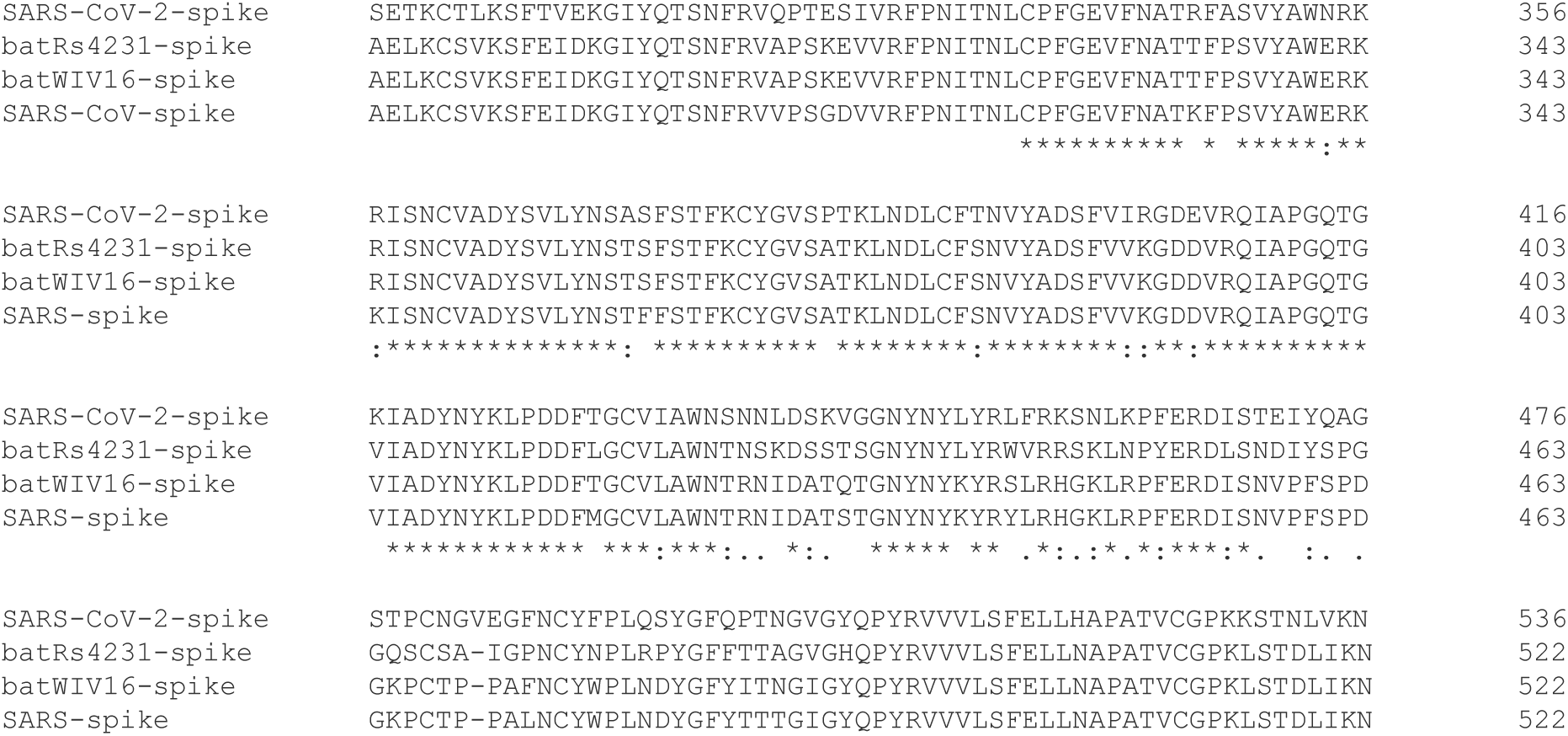
Sequence alignment of the receptor binding domain of SARS-CoV and related betacoronaviruses measured by Biolayer Interferometry for reactivity to SARS-CoV antibodies.

**Fig. S2.**
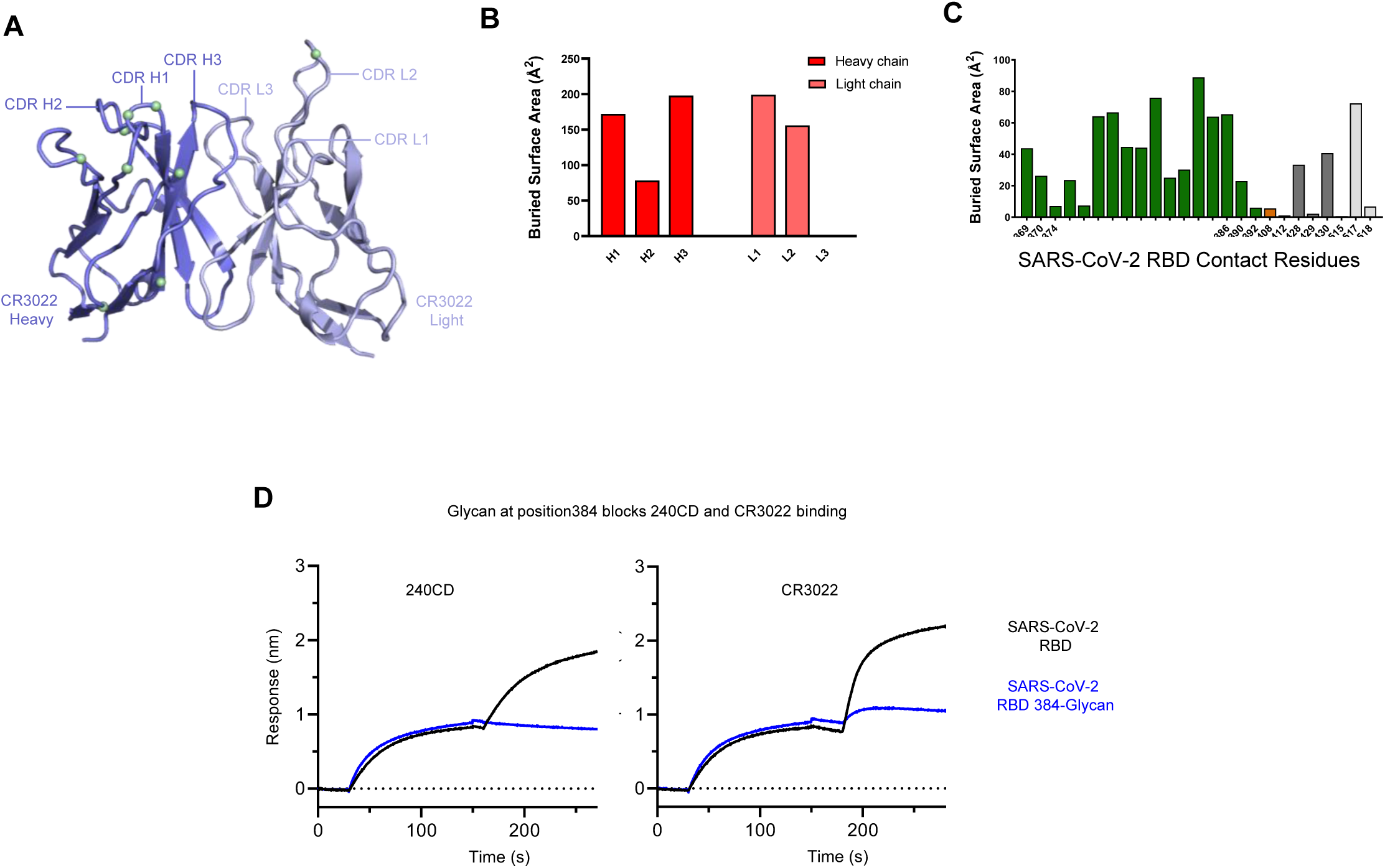
CR3022 Fab crystal structure and epitope analysis. **A** CR3022 is shown in ribbon representation, with CDR loops indicated. Residues which have undergone somatic hypermutation are indicated with green spheres. **B** Heavy and light chain CDR loops paratope buried surface area. **C** Buried surface area analysis of the CR3022 epitope on the SARS-CoV-2 RBD. **D** Antibody binding to SARS-CoV-2 RBD and RBD with a glycan at position 384. RBD is initially loaded onto the HIS probe, followed by incubation with RBD variants.

**Fig. S3.**
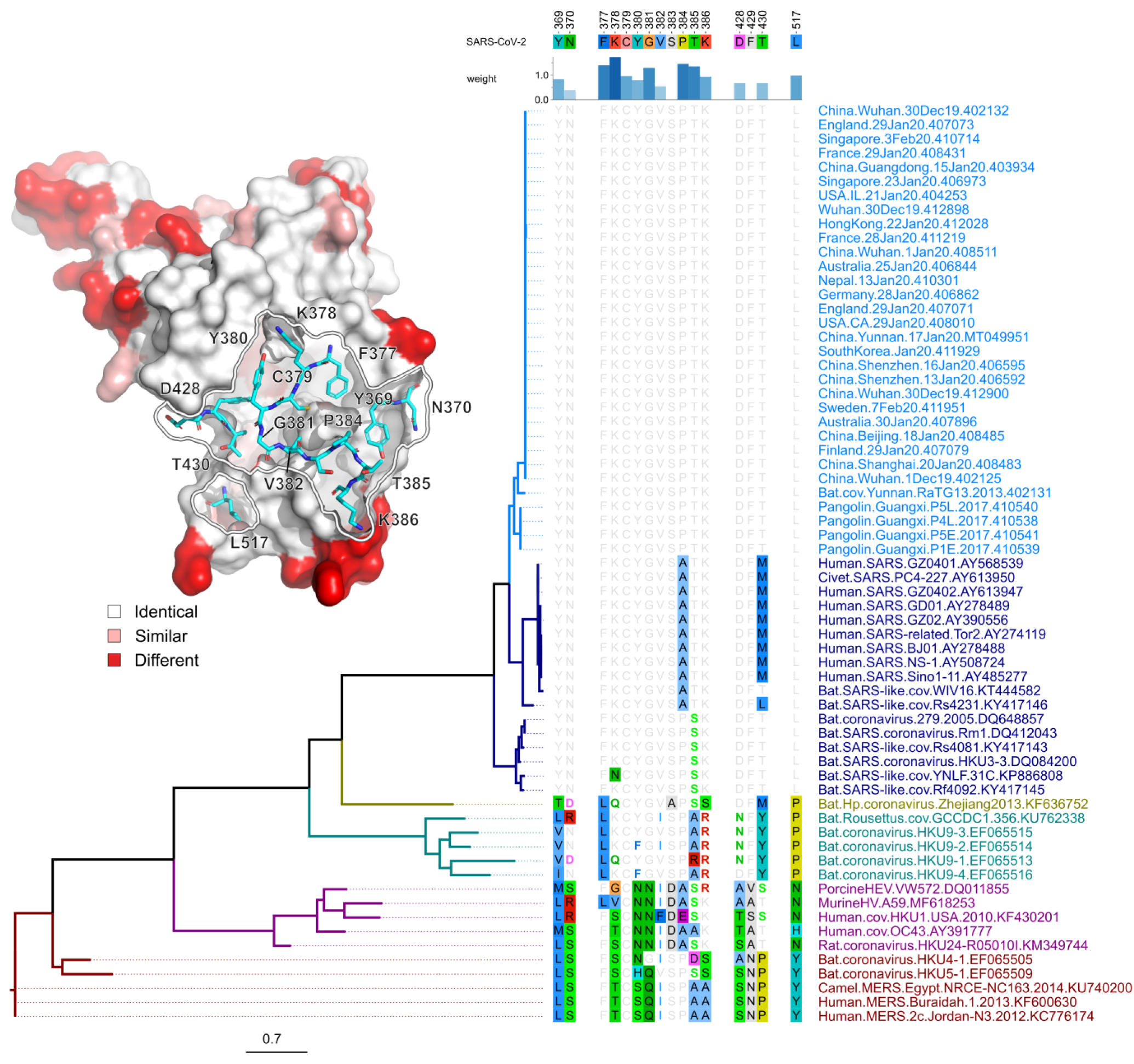
Structural and sequence analysis of the CR3022 epitope. Analysis of the CR3022 footprint across betacoronaviruses. The CR3022 epitope on SARS-CoV-2 (China.Wuhan.30Dec19.402132) RBD is compared across betacoronaviruses. The epitope is numbered according to the Wuhan reference; the strength of the interaction between the Ab and the spike protein is indicated by the height and color of the histogram bars above the sequence alignment. Sequences are ordered based on their phylogenetic relationships based on a maximum likelihood phylogenetic tree derived from amino acid RBD sequences. The RBD structure is shown in surface representation and depicts mutations between SARS-CoV-1 and SARS-CoV-2 in red; the CR3022 epitope is outlined in white, with contact residues shown in stick representation.

**Fig. S4.**
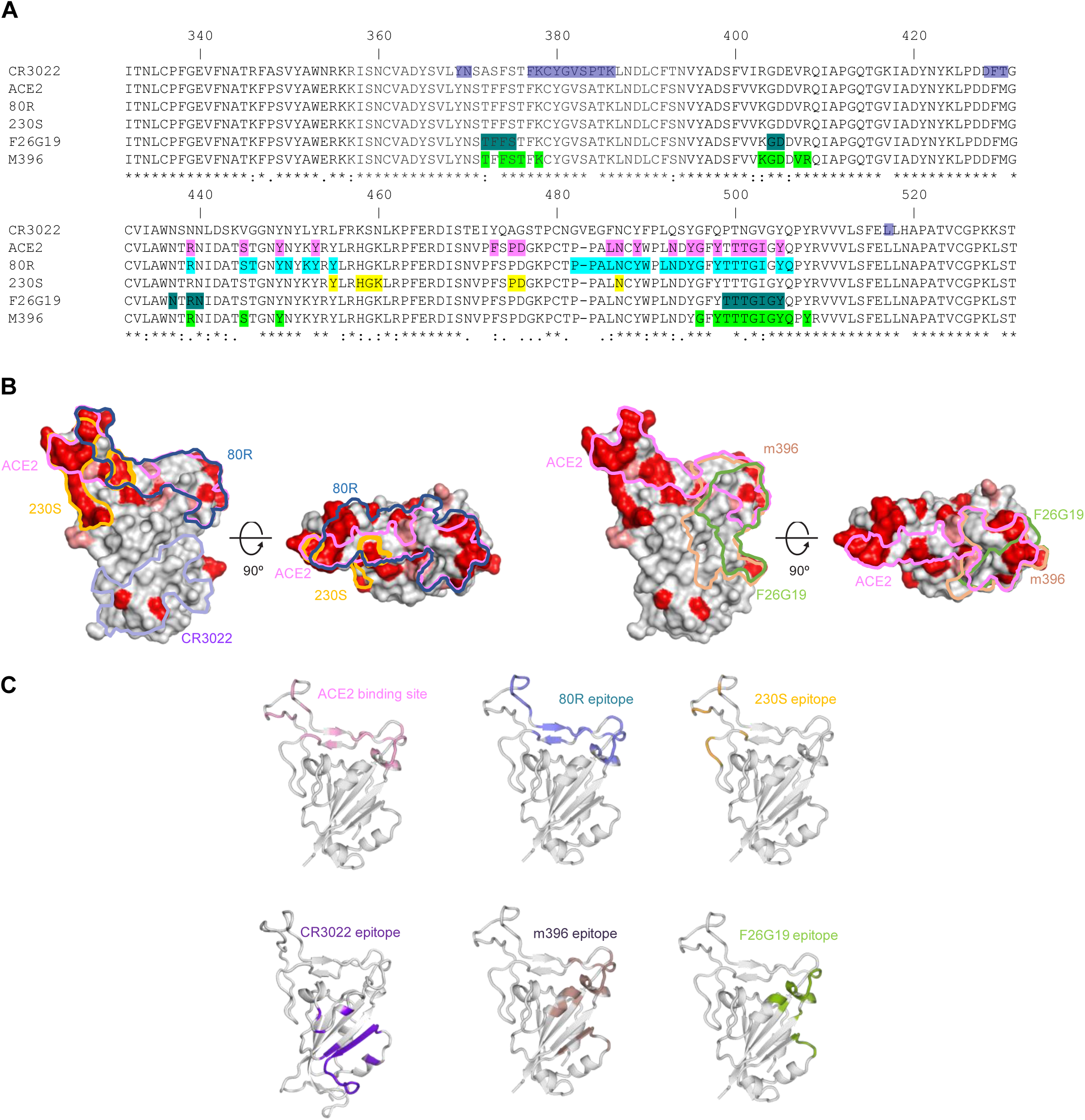
SARS-CoV-2 and SARS-CoV antibody epitope analysis. **A** Sequence alignment of SARS-CoV-2 and SARS-CoV RBD. Antibody and ACE2 receptor interacting residues are highlighted on the sequence. **B** Antibody epitopes and receptor binding sites are outlined on the surface of the SARS-CoV RBD structures. The Sequence differences between SARS-COV-2 and SARS-CoV are indicated by red coloring on the RBD surface. **C** Antibody epitopes are shown on the SARS-CoV RBD molecule depicted in ribbon representation. The CR3022 epitope is shown on the SARS-CoV-2 RBD molecule.

